# MxlPy - Python Package for Mechanistic Learning in Life Science

**DOI:** 10.1101/2025.05.06.652335

**Authors:** Marvin van Aalst, Tim Nies, Tobias Pfennig, Anna Matuszyńska

## Abstract

**Summary:** Recent advances in artificial intelligence have accelerated the adoption of ML in biology, enabling powerful predictive models across diverse applications. However, in scientific research, the need for interpretability and mechanistic insight remains crucial. To address this, we introduce MxlPy, a Python package that combines mechanistic modelling with ML to deliver explainable, data-informed solutions. MxlPy facilitates mechanistic learning, an emerging approach that integrates the transparency of mathematical models with the flexibility of data-driven methods. By streamlining tasks such as data integration, model formulation, output analysis, and surrogate modelling, MxlPy enhances the modelling experience without sacrificing interpretability. Designed for both computational biologists and interdisciplinary researchers, it supports the development of accurate, efficient, and explainable models, making it a valuable tool for advancing bioinformatics, systems biology, and biomedical research.

**Availability:** MxlPy source code is freely available at https://github.com/Computational-Biology-Aachen/MxlPy. The full documentation with features and examples can be found here https://computational-biology-aachen.github.io/MxlPy.

## Introduction

Mathematical modelling has evolved into a cornerstone of modern biological research, offering a rigorous, quantitative framework to explore the complexity of living systems [1]. Far beyond tracking metabolite concentrations, computational models enable the formulation and testing of hypotheses [1], prediction of emergent system behavior [2], and identification of control points in regulatory networks [3, 4]. They support the design of synthetic pathways [5], or even whole communities [6, 7], simulate therapeutic interventions [8, 9], and unravel nonlinear dynamics that are often inaccessible through experimental methods alone. Crucially, models serve as *in silico* laboratories, allowing researchers to conduct virtual experiments that are impractical or impossible *in vivo* or *in vitro* — whether due to cost, ethical constraints, or technical limitations. To enable the systematic construction and analysis of such dynamic biological models, we previously developed the Python package modelbase [10, 11], a toolkit tailored for building and simulating mechanistic models based on ordinary differential equations (ODEs). This package has been successfully utilized in a range of studies, particularly developing models of primary metabolism and photosynthesis [12, 13, 14, 15, 16], providing insights that complement and guide experimental efforts. However, the ongoing AI revolution is reshaping the computational biology landscape, where small-scale mechanistic models are increasingly being displaced by data-driven predictive approaches. In line with the perspective of Baker *et al*. [17], we believe that life sciences should embrace the complementary strengths of both mechanistic and machine learning (ML) methods. Motivated by this vision, we decided to expand our software to support hybrid modelling approaches, in particular mechanistic learning, that combine interpretability with predictive power.

### What is mechanistic learning?

**Mechanistic learning** (MxL) is an emerging approach that combines two fundamentally distinct modelling paradigms: mechanistic modelling and machine learning [18]. **Mechanistic models** - often implemented as systems of ODEs - are constructed based on known or hypothesized chemical, physical, and biological principles governing a natural process [19]. These equations capture how system variables evolve over time and are parameterized by quantities such as reaction rates, affinities, or transport constants. As models grow in complexity, the number of parameters can easily reach dozens or even hundreds. Despite efforts to systematically catalog measured parameter values in databases like BRENDA [20], SABIO-RK [21], and Bionumbers [22], such data are often sparse, inconsistent, or context-dependent. A major challenge arises from the fact that even when parameter values are available, they often reflect specific experimental conditions, developmental stages, or physiological states— rarely matching those under investigation in a given model. This introduces significant **uncertainty**, which can compromise the reliability of predictions.

To address these issues, researchers have developed techniques such as **ensemble modelling** (e.g., [23]), where plausible parameter sets are sampled from predefined statistical distributions and analyzed to assess the robustness of model behavior. However, parameter uncertainty is only one limitation of mechanistic models. Increasingly, approaches are needed that can deal with incomplete biological knowledge, guide model construction, and integrate experimental data more flexibly.

Rather than treating data-driven and knowledge-driven approaches as separate tools, mechanistic learning integrates them to create models that are both interpretable and adaptable to complex, uncertain systems. ML uses advanced statistical methods to extract information from data to make statements about a phenomenon, even if biological knowledge about a process is lacking, allowing accurate and fast predictions. However, a large amount of data must be acquired to train the underlying statistical model. Data acquisition can be a tedious task that, e.g., needs manual input for labelling. Many ML approaches are black boxes for which it is unclear how connections between input and output data are established during the training. Moreover, ML techniques are prone to overfit the data, thus forcing researchers to pay special care to robustness and generalizability.

Mechanistic learning combines the strengths of both approaches: using ML to infer unknown relationships, calibrate models, or augment incomplete mechanisms, while grounding model behavior in the structure and **interpretability** of mechanistic equations. While the term is still evolving, we follow the definition of [18], viewing mechanistic learning as the integration of ML and mechanistic modelling to study complex biological systems under uncertainty. This idea shares conceptual ground with related fields: scientific ML (e.g., [24, 25, 26]), used predominantly in physics and engineering, emphasizes embedding physical laws into ML models; or hybrid modelling more loosely refers to any combination of mechanistic and data-driven approaches [27]. What distinguishes mechanistic learning, especially in the life sciences, is the deliberate integration of ML into mechanistic modeling pipelines to complement and inform the model structure itself. In their review, Metzcar *et al*. distinguish four types of mechanistic learning [18]: sequential, parallel, extrinsic, and intrinsic. In the sequential type, the output of either a kinetic model or a ML approach is fed into the other as input. For instance, researchers developed neural networks to estimate parameter values for enzyme-catalysed reactions, which could be used in a mechanistic model (see [28]). Parallel mechanistic learning treats ML and mechanistic modelling as equivalent. Here, neural networks can serve as surrogate models that can replicate input-output relationships similar to mechanistic models without a deeper understanding of the underlying biology. Using surrogates allows researchers to make faster predictions by evading computationally costly differential equation integration [29]. The extrinsic approach employs mechanistic modelling for understanding post-hoc what neural networks have learned. Finally, intrinsic mechanistic learning uses domain knowledge in the development of statistical models (e.g., physics-informed (PINN)[25] or biology-informed neural networks (BINN) [30]). Overall, mechanistic learning can be regarded as an approach to constructing digital twins or multi-scale models ([31, 32, 33, 34]).

Despite its promise, mechanistic learning remains largely inaccessible to beginners and researchers without a strong computational background. Currently, few software tools support seamless integration of neural networks into mechanistic models. In most cases, such integration is implemented *ad hoc* within individual projects, without reusable, user-friendly infrastructure, e.g. [35, 36, 37] (with some notable exceptions, discussed in the supplement Table S1 [38, 39, 40, 41, 42, 43]).

To bridge this gap, we developed MxlPy — an enhanced and extended version of our previous toolkit, modelbase. MxlPy introduces both sequential and parallel mechanistic learning workflows, specifically designed to address the common challenge of missing or uncertain information in mechanistic models. With a clean, Pythonic interface, MxlPy empowers not only experts but also students and experimentalists to perform neural posterior estimation, assess parameter uncertainty using Monte Carlo (MC) methods, integrate surrogate models, or retrieve biologically relevant parameter distributions from open-access databases (see Fig. 1). By lowering technical barriers and embedding powerful inference tools, MxlPy makes mechanistic learning truly approachable for the broader life science community.

**Fig. 1.**
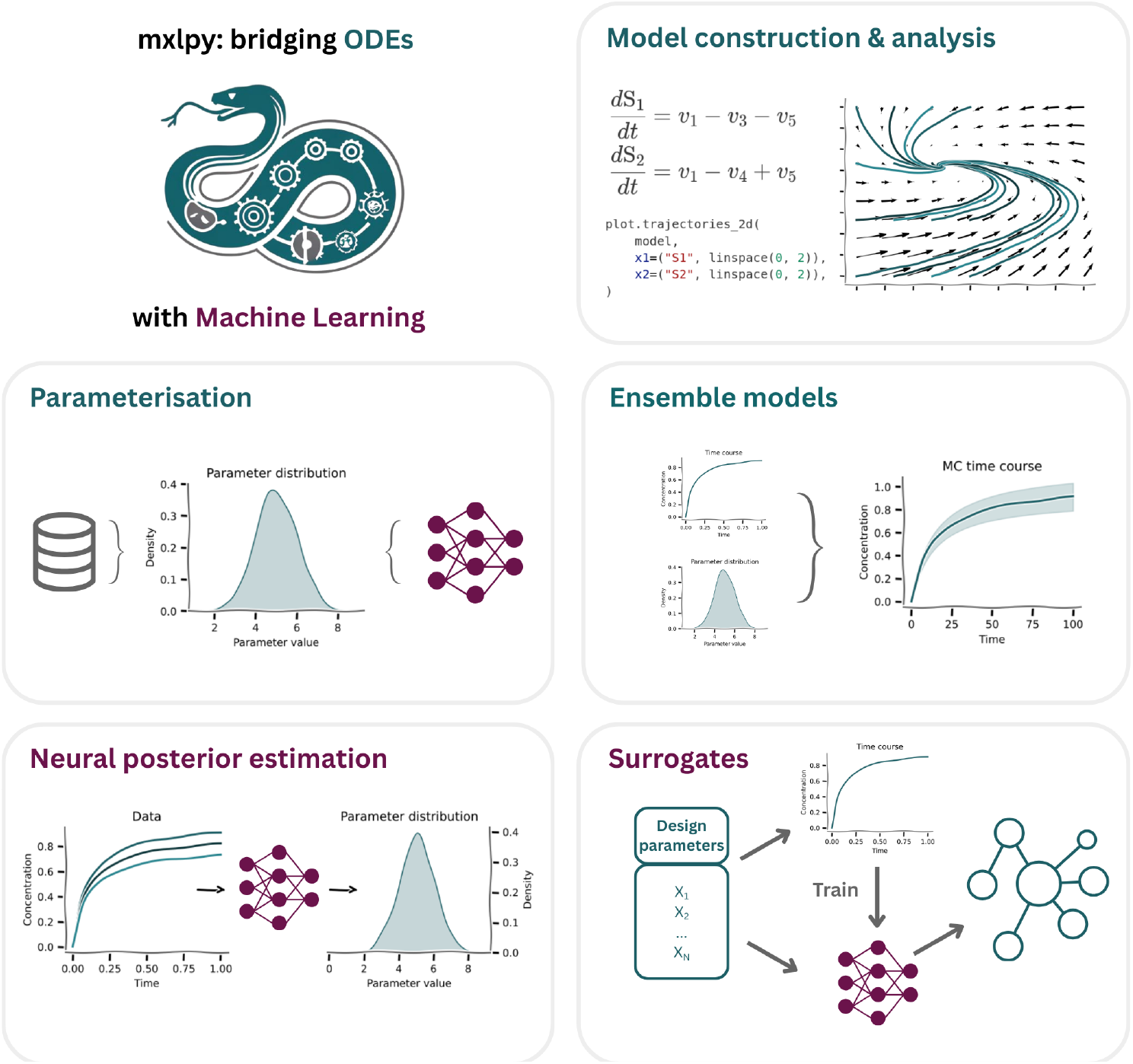
Key features of MxlPy enabling integrative kinetic modeling and machine learning. MxlPy facilitates mechanistic learning by combining traditional kinetic modeling with modern machine learning techniques. The platform supports direct parameter extraction from databases such as BRENDA, ensemble modeling for uncertainty quantification, and integration of surrogate models and neural posterior estimation methods. Designed for both experts and non-specialists, MxlPy streamlines parameter management, automates Monte Carlo simulations, and enables the hybridization of data-driven and mechanistic approaches for systems biology applications.

## Methods and Results

We present the core functionalities of MxlPy, a Python-based framework designed to integrate ML techniques with mechanistic modelling in the life sciences. MxlPy extends the capabilities of its predecessor, modelbase [11], by addressing a major limitation of mechanistic models: the scarcity and/or uncertainty of biological parameters. The current implementation of MxlPy provides five primary modules: mechanistic model construction and analysis, ensemble modelling (parameter sampling), surrogate model integration, parameter assessment from databases, and neural posterior estimation (Fig. 1).

In the following MxlPy’s facilities will be explained and showcased using examples. In particular, we will pay attention to those parts of the software that illustrate its potential for integrating ML, MC simulation, and uncertainty quantification into the mechanistic modelling process. The examples and analyses will be conducted using well-established, already-published models or adaptions of them. A comprehensive and regularly updated technical guide, including the reproducible examples, is available via the official MxlPy documentation portal: computational-biology-aachen.github.io/modelbase2.

### Modularised construction and analysis of kinetic models

Kinetic models in MxlPy are built programmatically using plain Python functions. The model objects provide full introspection and other meta-programming capabilities. These functionalities allow models to be built interactively and re-use parts of or the whole model in different computational projects, enabling modularisation via composition and inheritance. The key building blocks of kinetic mechanistic models (Eq. (1)) are parameters *ϑ*, variables *S*, and reactions specified by rate laws *v*(*t, ϑ*) (*t* is simulation time) and stoichiometries *N* (stoichiometric matrix),

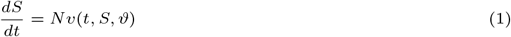

These building blocks and the simulation time can be used to derive additional stoichiometries, parameters, and variables. The type (static or dynamic) of derived quantities is determined automatically using the programmatic concept of lazy evaluation. Easy-to-use methods guide users through the specification of all parts of a computational model, thus allowing a gentle learning experience.

Time courses of dynamic variables can be simulated using various functions, including protocoled time courses that change a parameter during simulation according to a pre-defined switching protocol. The underlying ordinary differential equations of models are integrated using solvers implemented in the Python packages SciPy (e.g., LSODA) or Assimulo (CVode) [44, 45]. Users can fully customize all solver parameters, thus handling dynamically challenging integrations (e.g., stiff problems).

Several advanced functionalities are implemented in MxlPy, expanding upon its predecessor, modelbase [11]. Like modelbase, MxlPy supports steady-state analysis of biological systems through metabolic control analysis [46, 47]. Additionally, a LaTeX export feature is provided to facilitate the streamlined reporting of model structure by program introspection, providing ready to paste systems of equations for scientific publication, theses and other technical documents. A novelty in MxlPy is the automatic transformation of dynamic models into symbolic form using the SymPy library [48]. This symbolic representation enables integration with advanced mathematical frameworks for structural analysis of biochemical networks, including stoichiometric analysis, linear stability analysis, and **structural identifiability**. Structural identifiability analysis determines whether the parameters of a model can be uniquely estimated from noise-free, continuous observations, given the structure of the differential equations alone. Parameters that are structurally unidentifiable are inherently non-estimable regardless of the data quality or quantity [49, 50]. Detecting such parameters is essential in systems biology, particularly for interdisciplinary projects where precise parameter estimation underpins experimental validation and mechanistic insight. MxlPy integrates StrikePy, currently the only fully Python-based tool for structural identifiability analysis [51]. StrikePy uses symbolic computation to evaluate identifiability in small to moderately sized ODE models, thus complementing MxlPy’s symbolic infrastructure.

### Parametrisation

MxlPy provides user-friendly methods for extracting and integrating values of kinetic parameters from the BRENDA enzyme database [20] into models. A copy of an enzyme’s entry or a set of entries must be downloaded from the database as a JSON file. Functionalities in the MxlPy’s parameterise module help extract the desired information from the downloaded database entries and provide the user with a data frame that facilitates further analysis.

### Ensemble modelling and Monte Carlo sampling

MxlPy provides a dedicated framework for ensamble simulation using Monte Carlo (MC) sampling through its mc and distributions modules. Users define the uncertainty in each model parameter by specifying appropriate statistical distributions (among others, uniform, normal, log-normal), e.g.,

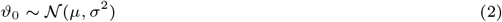

These distributions can be derived from domain expertise, experimental measurement ranges, or extracted directly from biological databases (e.g., BRENDA) when multiple empirical values are available for a given parameter across organisms or conditions. Once the parameter distributions are specified, MxlPy performs repeated sampling to generate a user-defined number of parameter sets. Each set is used to simulate the model, producing an ensemble of **time course trajectories** *S*_*i*,*t*_. These can be statistically summarized

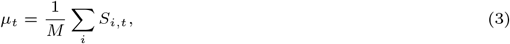

where *M* is the number of dynamic variables, or visualized using built-in plotting functions to evaluate system robustness, identify sensitive parameters, or quantify variability in predicted behaviors. In addition to dynamic simulations, MxlPy supports MC-based analysis of protocol-driven simulations (e.g., experimental perturbation protocols), steady-state properties, and metabolic control coefficients. All simulations are executed via a unified interface (e.g., mc.time course), allowing for reproducible and scalable ensemble workflows with minimal overhead.

The usage of the ensemble model functionalities will be showcased with the following example.

Photosynthesis is an important part of the complex biological, chemical, and physical network that sustains life on Earth. Next to specialized microbial life, plants are the organisms that show photosynthetic activity, thereby fixing radial solar energy into chemical forms. The captured energy is used to fixate carbon dioxide into other organic compounds that are the building blocks for biomass. The Calvin-Benson-Bassham cycle (CBB) performs the carbon dioxide fixation, while the energy transformation happens in the photosynthetic electron transport chain (PETC). Both processes are located in the chloroplasts of leaves. Additionally, the electron transport through the PETC ultimately leads to oxygen release. The gates that mediate the gas (carbon dioxide and oxygen) exchange between leaves and the environment are stomata, small openings in the epidermis that open and close based on various environmental or physiological cues.

Kirschbaum *et al*. [52] developed a semi-empirical mechanistic model of stomata opening in response to light intensity in the plant *Alocasia macrorrhiza*. By simulations and parameter variations, Kirschbaum et al. could show that their model performs reasonably in changing light conditions. In particular, its capability to replicate transient responses of stomatal conductance *g*_*s*_ was promising and distinguished it from other modelling approaches in the late 1980s. The mathematical background of the model consists of three ordinary differential equations (ODEs) representing a not-specified biochemical signal *S* that is directly influenced by the applied light intensity, the osmotic potential *π* in the guard cells surrounding the stomata, and the leaf water potential *w. g*_*s*_ is proportional to *w*. All three ODEs follow simple first-order kinetic, each of which is specified by a rate constant determining the change of the biochemical signal (*τ*_*i*_, *τ*_*d*_, for increasing and decreasing signal), the osmotic (*τ*_*π*_) and the water potential (*τ*_*w*_).

As Kirschbaum *et al*. [52] state in their discussion, the rate constants of the processes determining the stomata conductance could vary depending on the leaf’s physiological state. Here, we leverage MxlPy’s MC and parallel simulation facilities to conduct a time-dependent global sensitivity analysis by combining it with the sampling and analysis methods provided by the package SAlib [53, 54]. With this, we try to answer which parameter variation significantly influences the stomatal conductance in a changing light environment. A global sensitivity analysis consists of several steps. Firstly, we choose the parameters *τ*_*i*_, *τ*_*d*_, *τ*_*π*_, *τ*_*w*_ as input that is varied and the time-course of the stomatal conductance *g*_*s*_ as output that is analyzed. Secondly, for the input parameters, we assume Gaussian priors with mean to the value given by Kirschbaum *et al*. and variation such that 99.7 % of the drawn parameters are located between 80 % and 120 % of the original values. Thirdly, to sample the parameter space evenly, we choose Sobol sequences that minimize discrepancy (not sampled regions) in the parameter space. Lastly, we decided to use Sobol sensitivity analysis, a variance-based technique indicating what percentage of the output variance is determined by the input variance. The Sobol method provides three different metrics. While the first-order sensitivity *S*_1_ indicates the influence of variance in a specific parameter on the output, the second-order sensitivity *S*_2_ specifies the combined effects of parameter variations. The total-order sensitivity combines both first- and second-order effects into one metric. Discrepancies between first- and total-order sensitivity are a good hint for higher-order parameter interactions determining changes in the output.

Fig. 2 shows the first-order sensitivity for variations of the input parameters during the application of two consecutive 5-min light pulses `a 500 μmol m^−2^ s^−1^ interrupted by dark phases to a previously dark-adapted leaf. Also represented is the average time course of the stomatal conductance *g*_*s*_. After applying a light pulse, the conductance has a delayed reaction. It reaches its maximum in the intermittent dark phases, followed by its return to the dark-adapted original state. As is visible by the colored areas around the *g*_*s*_ average trajectory, the chosen variation in the kinetic constants did not influence the time course drastically.

**Fig. 2.**
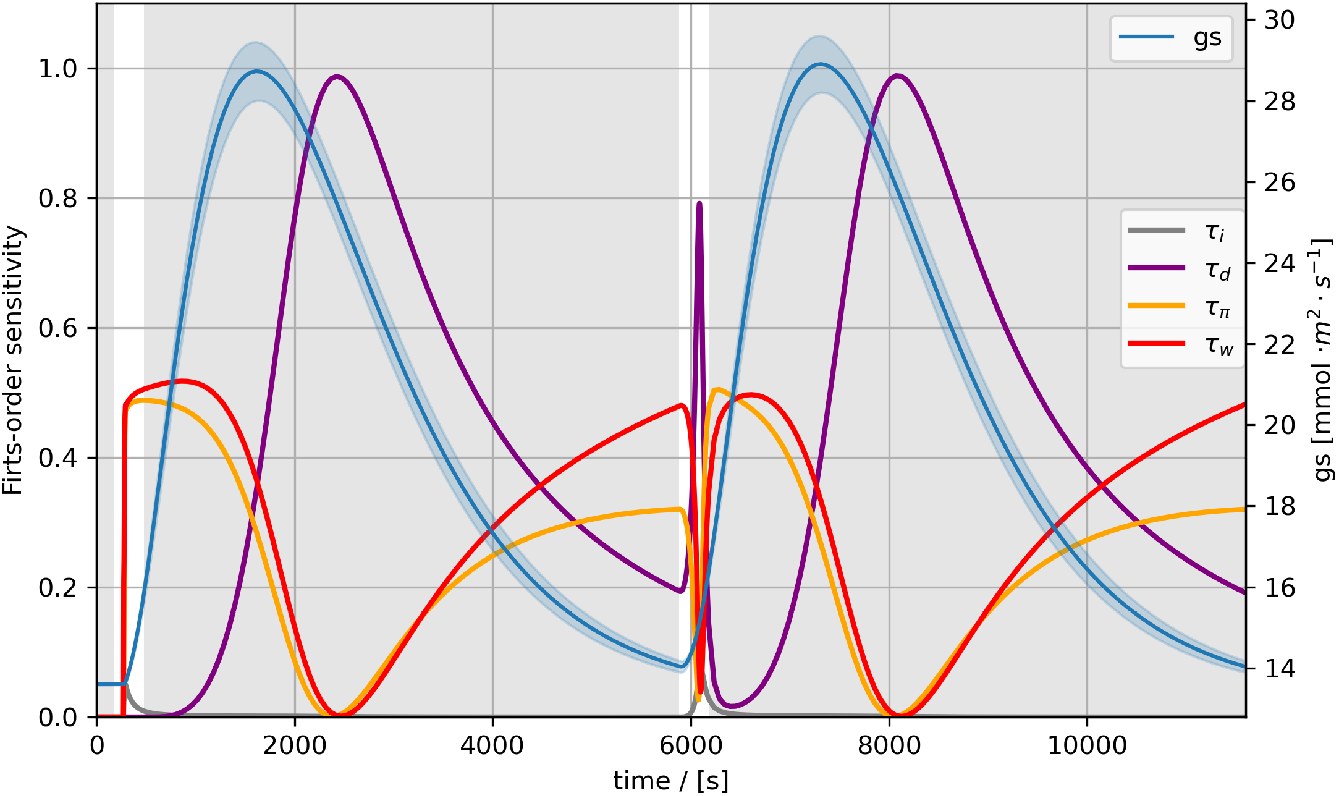
Global sensitivity analysis of the stomata conductance by Kirschbaum *et al*. [52]. First-order sensitivities *S*_1_ (purple, red, orange, and grey line) and average trajectory of the stomatal conductance *g*_*s*_ (blue line). Two consecutive 5 min light pulses `a 500 μmol m^−2^ s^−1^ interrupted by dark phases applied to a previously dark-adapted leaf are shown. The parameters varied for the sensitivity analysis were the rate constant for the biochemical signal, the osmotic, and the water potential in the leaf (*τ*_*i*_, *τ*_*d*_, *τ*_*π*_, *τ*_*w*_).

By the Sobol sensitivity analysis, we could explore which of the rate constant variations had the most effect at which time point during the experiment. Fig. 2 shows that, from the onset of a light pulse to the dark phase, the rate constants of water and osmotic potential are responsible for the changes in the *g*_*s*_ curve. Only later, the biochemical signal *τ*_*d*_ rate constant takes over and dominates the variations in *g*_*s*_ shortly after its maximum. Interestingly, the regions of the highest variations in the *g*_*s*_ curve are characterized by high *S*1 sensitivities for all rate constants except *τ*_*i*_, which only plays a role during a light pulse (compare maximum and decline of *g*_*s*_ trajectory). A comparison with *S*_*T*_ sensitivities (not shown) indicates that parameters have little second-order effects on the variation of stomatal conductance.

The sensitivity analysis, hence, shows that especially the rate constant of the osmotic potential and the leaf water potential play a critical role in the onset and closing of stomota while the rate constant of the biochemical signal is more important once the stomata are fully open.

### Surrogate models integration

Besides missing parameter information, a lack of structural details (equations, network structure) significantly hinders building models. Especially for more advanced model projects with different compartments, physical, chemical or biological details are often not fully known. However, gap-free computational representations of natural processes are necessary for making precise predictions informing, e.g., medical decision-making or creation of digital twins [55, 56, 57, 58]. Surrogate models that utilise ML methods have been established to address information gaps by adding statistical models,

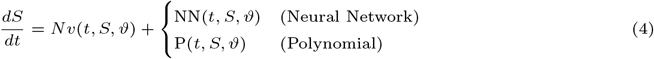

Moreover, surrogate models can replace mechanistic model parts in intensive simulations due to their lower computational cost after training.

MxlPy allows users to train surrogates and integrate them into models. Two surrogate approaches are implemented: neural networks and polynomial functions. MxlPy provides a user-friendly method for training neural network surrogates with PyTorch, train torch surrogate. Using training data, for instance, concentrations of substrates and metabolic fluxes, a neural network model is trained for a specified number of epochs. A multi-layer perceptron (input layer, a hidden layer with 50 neurons/ReLU and an output layer) is used by default. However, users can also provide custom PyTorch neural networks. Train types (full batch, mini-batch) and hyperparameters are customisable. Once a surrogate is trained, MxlPy allows the integration of surrogates in an existing model with the add surrogate method of a model object. Polynomial surrogates are implemented via the train polynomial surrogate function. By default, this function trains various polynomials up to the seventh degree. The degrees can be customized by users.

Surrogates could, for example, be used in photosynthetic organism models simulating anabolic pathways, e.g., chlorophyll synthesis. Surrogates trained on detailed models of light reactions (e.g. [12, 15]) would allow for computationally cheap estimations of reactant productions, fueling the anabolic model, see Fig. 3. This would particularly improve performance when the models differ in time resolution. Beyond computational efficiency, such surrogate integration enables biotechnological applications, such as the rational design of strains optimized for metabolite production under specific light conditions. By tailoring models based on the availability of energy carriers like ATP and NADPH, researchers can better predict and engineer productivity in photosynthetic systems.

**Fig. 3.**
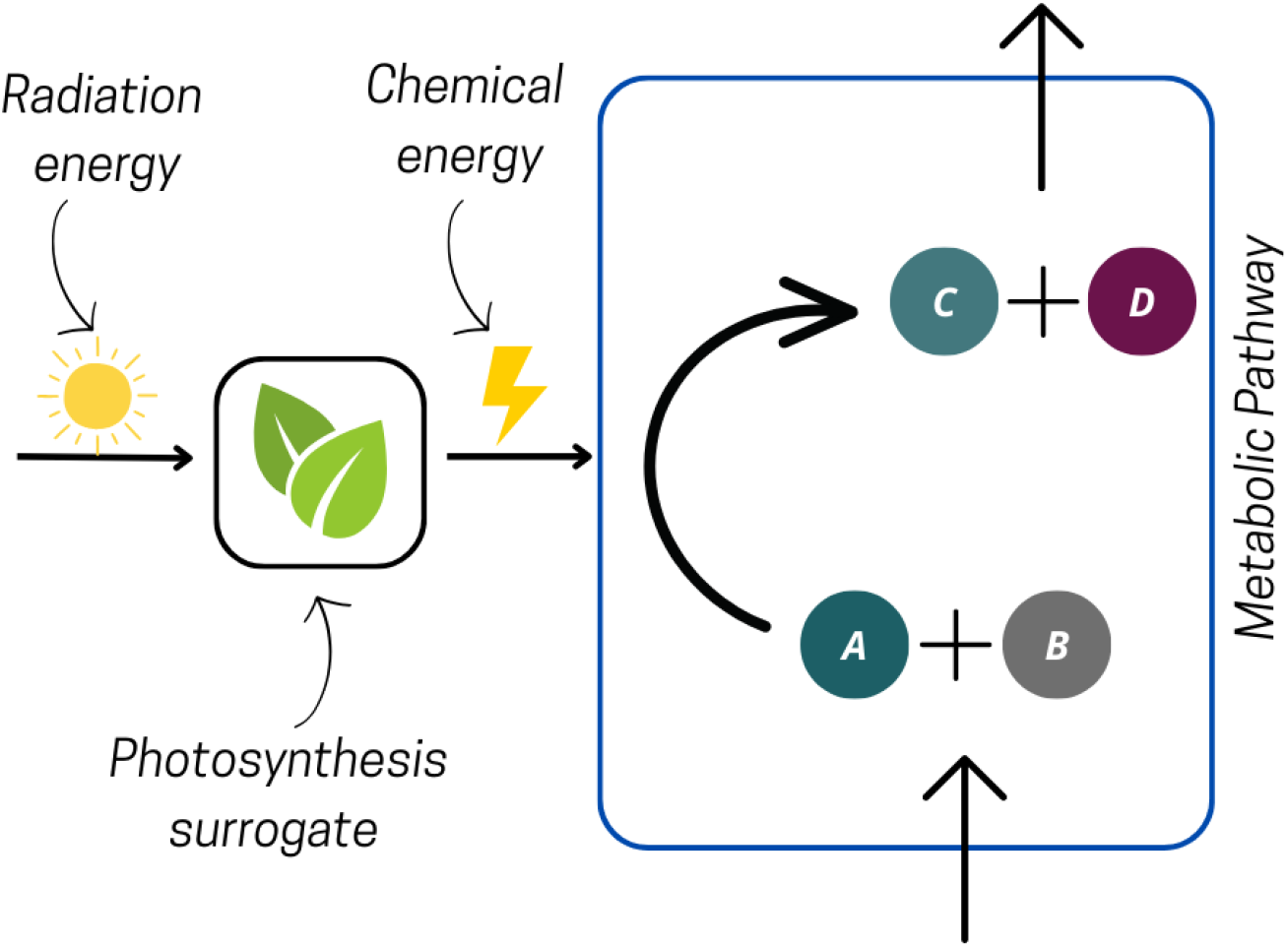
Scheme of a photosynthesis surrogate. The details of the photosynthetic electron transport chain have been black-boxed using ML techniques. This allows for a computational efficient way to integrate the light reactions into mechanistic models of secondary metabolism in photosynthetic organisms, maintaining complex responses to varying light environments.

### Neural posterior prediction

Neural posterior estimation (NPE) is a technique that, given a model structure and parameter priors, learns to predict the parameter posterior distribution. The frequent intractability of the data likelihood function in model studies, which is needed for classical Bayesian approaches, motivated researchers to develop computational advanced simulation-based inference techniques (SBI) consisting of Approximate Bayesian Computation and Neural density estimation [59, 60, 61, 62, 63, 64]. NPE circumvents the regular Bayesian workflow of collecting priors, deriving a likelihood function, and calculating the posterior via the Bayesian formula, using a neural network instead. This neural network directly predicts the posterior distributions of model parameters given the data. It is trained on a model-informed synthetic data set combined with parameter priors.

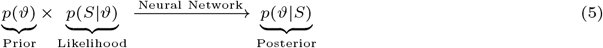

MxlPy provides an easy-to-use npe module that allows users to specify simulation-based inference, particularly NPE workflows. For this, the users must first specify a suitable prior distribution for the model parameters. The prior distribution can be derived from database information using MxlPy’s parameterisation functionalities or domain knowledge. A synthetic data set is created using the prior distributions by leveraging MxlPy’s scan and distributions modules, facilitating the sampling of parameters and simulations over a range of parameter values. Simulated data, such as time courses, steady states, control coefficients, or summary statistics, is then used to train a neural network that predicts a parameter set given a synthetic data sample. For this, the user can choose one of the npe module’s training functions, allowing access to PyTorch. The trained neural network can be used as an approximation for a posterior distribution function, describing the parameter probability for a given data set via density estimation. The default neural network in the npe module is the same as described for the surrogate module (see above), but can be changed by the users to use state-of-the-art neural network architectures presented in the SBI literature.

## Discussion and future prospects

Mathematical mechanistic models serve, among others, to create hypotheses and predict future events. These two purposes require the modeler to develop a precise and interpretable formalization of a biological process. However, precision is especially limited by the uncertainties about the model structure and its parameter values. An advanced computational tool that allows experts and non-experts to implement mathematical models should facilitate uncertainty quantification and help users mitigate the effects of unknown model properties. Mechanistic learning is well-suited for the latter because it enables the replacement of important but uncharacterized parts of a biological process with a statistical model. This statistical model captures the experimental input-output relationship of the uncharacterized part.

In this manuscript, we presented MxlPy, a Python package that enables expert and non-expert users to construct ODE-based models and analyze them with classical (not-shown but compare [11]) and mechanistic learning techniques. We introduced MxlPy’s five current core functionalities - mechanistic model construction and analysis, ensemble modelling (parameter sampling), surrogate model integration, parameter assessment from databases, and neural posterior estimation. The parameter access functionality allows users to efficiently process parameters from public databases, particularly BRENDA. MxlPy enables ensemble modelling via its parallel simulation options and MC functions. Together, these two functionalities provide the basis for the parallel exploration of a network structure with parameters drawn from prior distributions. These prior distributions can be obtained by the analyses of parameter database entries or specified by the users using domain knowledge. MxlPy allows the creation of surrogates based on datasets describing a model or model part’s inputs (parameters) and outputs (simulated state trajectories). Surrogates simplify an existing model or black-box an uncharacterized model part, thus mitigating model uncertainties or helping conduct extensive sensitivity analysis. MxlPy also enables users to perform neural posterior estimation, an advanced mechanistic learning technique that derives parameter probabilities based on domain knowledge encoded as a model.

We showed MxlPy core functionalities with examples and references. We implemented a mathematical description of stomata conductance in plant leaves. Stomata are small openings that allow gas exchange between the leaf and the environment. By combining MxlPy’s ensemble model function with the advanced global sensitivity methods provided by the package SAlib, we could investigate the uncertainties in the stomata conductance that arise due to changes in parameter values. This is especially critical since those parameters are assumed to vary based on the plant’s physiological status.

To our knowledge, MxlPy is, with its integration of ML at the forefront of software, providing users with mechanistic learning options in the modelling process. However, MxlPy also shares aspects with a series of other computational tools, allowing users to simulate life science systems, as summarised in Tab. S1. An example is the CADET modelling framework that emphasizes chromatography and biotechnological process modelling [43]. CADET provides options for binding, reaction, and population balance models. Moreover, MxlPy shares similarities with other classical system biology toolboxes, like Copasi, Tellurium, BioNetGen, or VCell [40, 42, 65]. However, the merit of MxlPy is its Pythonic, easy-to-use architecture specifically designed to implement ML and ensemble model approaches into mechanistic modelling projects. MxlPy’s structure facilitates systematic construction, replication, and analysis of models with and without mechanistic learning. No domain language is established to describe and formalize computational models, allowing users to employ their knowledge of Python. MxlPy’s architecture as standalone software distinguishes it from other Python software projects, such as MEMMAL [66], a pipeline combining ML and mechanistic modelling using several Jupyter notebooks. Additionally, MxlPy shares common ground with the HybridML project [38]. Both tools allow the integration of neural networks into the model process. However, MxlPy emphasizes but is not constrained to metabolic models. MxlPy allows, for instance, metabolic control analysis in ensemble modelling to identify critical steps in industrial or medical-relevant metabolic pathways with uncertain parameters. Taking look into other programming languages it is impossible not to mention the SciML ecosystem (Scientific ML in Julia) and its Catalyst.jl. This Julia-based software package for modeling and simulating reaction networks using symbolic representations of chemical or biochemical systems serves a similar role to the Julia community as MxlPy aims to do for the Python community [39]. Both software projects provide options to replace rate laws in mechanistic models with data-driven functions. While scientific ML broadly focuses on embedding physical or biological constraints into neural architectures, MxlPy was created to serve the specific and practical needs of life science modelers—offering a transparent, modular framework that supports end-to-end workflows for kinetic model construction, calibration, hybridization with ML components, and reproducible analysis within the context of complex experimental collaborations.

Software development is an evolving process, and MxlPy is no exception. To enhance its utility and robustness, we plan to expand MxlPy with additional modules aimed at better managing uncertainty and evaluating a model’s capacity to generate hypotheses and make reliable predictions. Future developments will focus particularly on quantifying parameter sloppiness and addressing model unidentifiability—challenges that are especially significant in collaborative efforts between computational and experimental researchers. In particular, the communication with existing packages for ML-based estimation of enzymatic properties (e.g., catalytic constants) will be supported [67]. These enhancements will support more precise parameter estimation, even in complex modeling scenarios. To foster transparency, reproducibility, and community-driven innovation (see Table S2 for a comparison with FAIR4RS principles [68]), MxlPy is available as an open-source project, and we warmly invite contributions and feedback from the broader scientific community.

## Conclusion

MxlPy is a Python package designed for user-friendly model creation and analysis. Following the mechanistic learning principle, it combines the interpretability of mechanistic models with the prediction capabilities of ML. It supports model creation, parameterization, and analysis using state-of-the-art methods, such as surrogate models, MC sampling, and neural posterior estimation. With MxlPy, we want to allow experts and non-experts alike to create reliable and replicable models.

## Abbreviations

BINN: biology-informed neural networks
MC: Monte Carlo
ML: machine learning
MxL: mechanistic learning
NPE: neural posterior estimation
ODE: ordinary differential equations
PINN: physics-informed neural networks
SBI: simulation based inference

## Data availability

The source code of MxlPy can be found on GitHub (github.com/Computational-Biology-Aachen/MxlPy). The documentation can be accessed at computational-biology-aachen.github.io/MxlPy.

## Competing interests

No competing interest is declared.

## Author contributions statement

**M.v.A**.: methodology; formal analysis; software resources; writing – original draft; writing – review & editing, **T.N**.: software resources, writing – original draft, formal analysis; writing – review & editing, **T.P**.: writing – original draft; writing – review & editing, **A.M**.: conceptualization and supervision, funding acquisition; writing – original draft, writing – review & editing

## Funding

The Deutsche Forschungsgemeinschaft (DFG) is acknowledged for financial support via Germany’s Excellence Strategy (EXC 2048/1, project number: 390686111), the DFG Research Unit FOR 5573 “Dynamic Regulation of the Proton Motive Force in Photosynthesis” (GoPMF, project number: 507704013), and SFB 1535: “Microbial networking – from organelles to cross-kingdom communities” (MibiNet, project number: 458090666).

## Supplementary information

### Software Comparison

The following table (Tab. S1) compares various software outlining the need for frameworks that allow easy integration of machine learning/neural networks into mechanistic models. We assumed a feature would be available if a function in the software allowed specific analyses (either as direct implementation or wrapper for third-party software). However, if just examples were given in the documentation of using third-party software directly, we defined a feature not available in the software itself. Stability analysis refers to linear stability analysis of complex systems. N/A indicates that we could not find any indicator that the feature is implemented. Although care was taken to find all relevant information in the documentation of each software, the reader is referred to each software’s website for detailed descriptions of its functionalities.

**Table S1.**
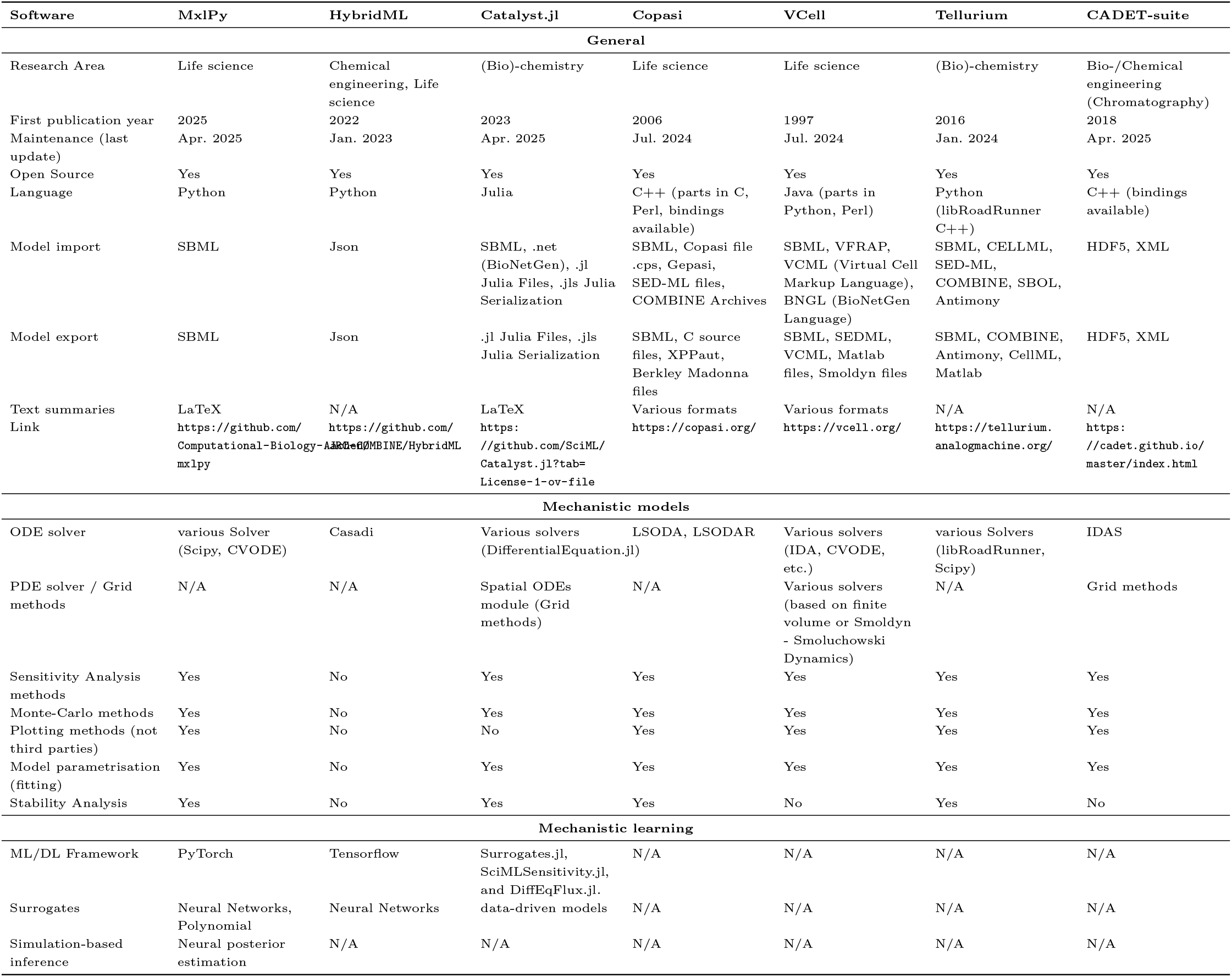
Comparison of modeling software platforms.

### FAIR4RS principles

Computational analyses of complex biological systems have become increasingly relevant with the advancement of data-intensive experiments. However, finding suitable, maintained and usable software can be challenging. To address these challenges, FAIR4RS [68] was introduced based on the FAIR (Findable, Accessible, Interoperable and Reusable) principles as part of the open science community. In the following section, we will compare how much MxLpy complies with the FAIR4RS principles.

**Table S2.** MxlPy alignment with FAIR4RS principles.

| Code | Principle Description | Compliance |
| --- | --- | --- |
| F | **Findability:** Software and metadata are easily discoverable by humans and machines. |  |
| F1 | Unique ID. | MxLpy has a unique PyPI (Python Package Index) name and a Zenodo identifier |
| F2 | Rich Metadata | Rich Metadata can be found in the Python Package Index or on GitHub |
| F3 | ID in Metadata | MxLpy's name is included in the metadata on PyPI. The Zenodo identifier is available on the GitHub page |
| F4 | Searchable Metadata | MxLpy's metadata can be searched |
| A | **Accessibility** |  |
| A1 | Retrieval Protocol | MxLpy can be downloaded via the browser on PyPI, following the links provided in the metadata using https or via pip. |
| A2 | Metadata Persistence | The metadata on PyPI is independent of the MxLpy repository and remain persistent once the repository becomes inaccessible |
| I | **Interoperability** |  |
| I1 | Standards Compliance | MxLpy uses standard formats for import and export of the system biology community (SBML) |
| I2 | Qualified References | MxLpy refers to other object, such as websites or links in documentation and code annotation |
| R | **Reusability** |  |
| R1 | Multiple Attributes | MxLpy is licensed under the GPL-3.0 licence. Detailed information and development history is accessible on the projects public GitHub page |
| R2 | Software References | The code includes detailed information about dependencies and allows installing dependencies manually via pixi.lock files. |
| R3 | Community Standards | MxLpy follows community standard in life science (system biology) community |

## Notes

https://github.com/Computational-Biology-Aachen/mxlpy

### Competing Interest Statement

The authors have declared no competing interest.

